# Engineered Bxb1 variants improve integrase activity and fidelity

**DOI:** 10.1101/2024.10.21.619419

**Authors:** Joshua Rose, Hyaeyeong Kim, Jessica Von Stetina, Mahant Thangavelu, Hasahn Conway, Mollie S. O’Hara, Patrick Hanna, Thomas Biondi, Maike Thamsen Dunyak, Jonathan D. Finn, Sandeep Kumar, Jesse C. Cochrane

## Abstract

Many current genome editing technologies rely on the action of large serine integrases (LSIs) to insert gene-sized DNA sequences into the genome. Bxb1 is the most commonly used LSI for therapeutic efforts, including PASTE, PASSIGE and I-PGI. While Bxb1 demonstrated good activity *in vitro* in cycling cells, the activity in non-dividing hepatocytes was significantly less efficient. Further, wild-type Bxb1 is known to have detectable off-target activity at cryptic genomic sites, which presents a potential safety risk for therapeutic development. To address these issues, we developed Bxb1 variants that demonstrate increased specificity and potency *in vitro* and have engineered stabilized Bxb1 variants that increase *in vivo* activity over 25-fold enabling targeted integration at therapeutically relevant levels.

---

Large serine integrases (LSIs) can insert any size and sequence of DNA into a specific location, given the presence of two complimentary binding sites for the LSI (1). This has inspired both gene therapy applications as well as cell line creation technologies (2-9). All LSIs contain four major structural domains, two of which, the recombinase and zinc ribbon, are responsible for recognizing the LSI’s DNA binding sites (10). Following binding of the DNA substrates (attB and attP), LSIs undergo a structural rearrangement and create a new DNA topology resulting in the insertion of DNA (11).

For many applications, the LSIs that seem to be the most studied are Bxb1 and PhiC31 (12). Bxb1 appears to be more active overall and has emerged as the enzyme of choice for large gene insertion therapies (6,8,9,13). Bxb1 can insert large DNA sequences without relying on any endogenous cellular DNA repair machinery (14). The cargo is delivered as a fully synthesized gene without need for DNA repair as the insertion is done seamlessly between two short attachment site sequences (10,11).

However, Bxb1 has two previously characterized limitations: specificity (off-targets) and low activity in certain cell types (6,7,13,15). Bxb1 can use imperfect attachment sites (cryptic sites) as locations for integration, several of which are present in the human genome (7,9,15). Wild-type Bxb1 also has relatively modest activity in cells (8,16). This makes it impractical to use for gene insertion as most cures rely on a minimum amount of active protein expression to be viable.

Here we describe a set of engineered and naturally occurring Bxb1 variants that overcome these limitations. These variants introduce mutations that lead to highly specific integrases and stabilization tags that increase *in vivo* activity. We anticipate that these enzymes will be of significant use in further efforts for large gene insertion for therapeutic benefit.

## Results

### Bxb1 is a highly active, site-specific recombinase

Four molecules of a large serine integrase (LSI) bind two DNA substrates, attB and attP, and catalyze a topological rearrangement leading to two DNA products, attL and attR (Fig. 1a) (11). LSIs have a conserved structural organization: the N-terminal domain contains a dimerization domain and the catalytic serine, two domains for DNA binding, a recombinase domain, and a zinc ribbon domain and a coiled-coil domain that mediate tetramer formation between two LSI dimers (1). For Bxb1, the DNA sequences of the substrates and products are well-defined and contain known binding sites for the recombinase and zinc ribbon domains (17). In the absence of DNA, Bxb1 is a homodimer (Supp. Fig. 1a). Using gel shift assays with fluorescently labeled substrates, we can accurately measure the affinity of Bxb1 for DNA substrates (Fig. 1b). Bxb1 dimers bind with relatively high affinity to both the substrate and product with Bxb1 dimers binding to the product sequences, attL and attR, with affinities of around 3 nM, and to the substrate sequences, attB and attP, with slightly weaker affinity of around 12 nM (Fig. 1c). At high concentrations, slow migrating species forms, suggestive of non-productive multimerization, but there is no evidence for DNA rearrangement without both substrates being present.

**Figure 1.**
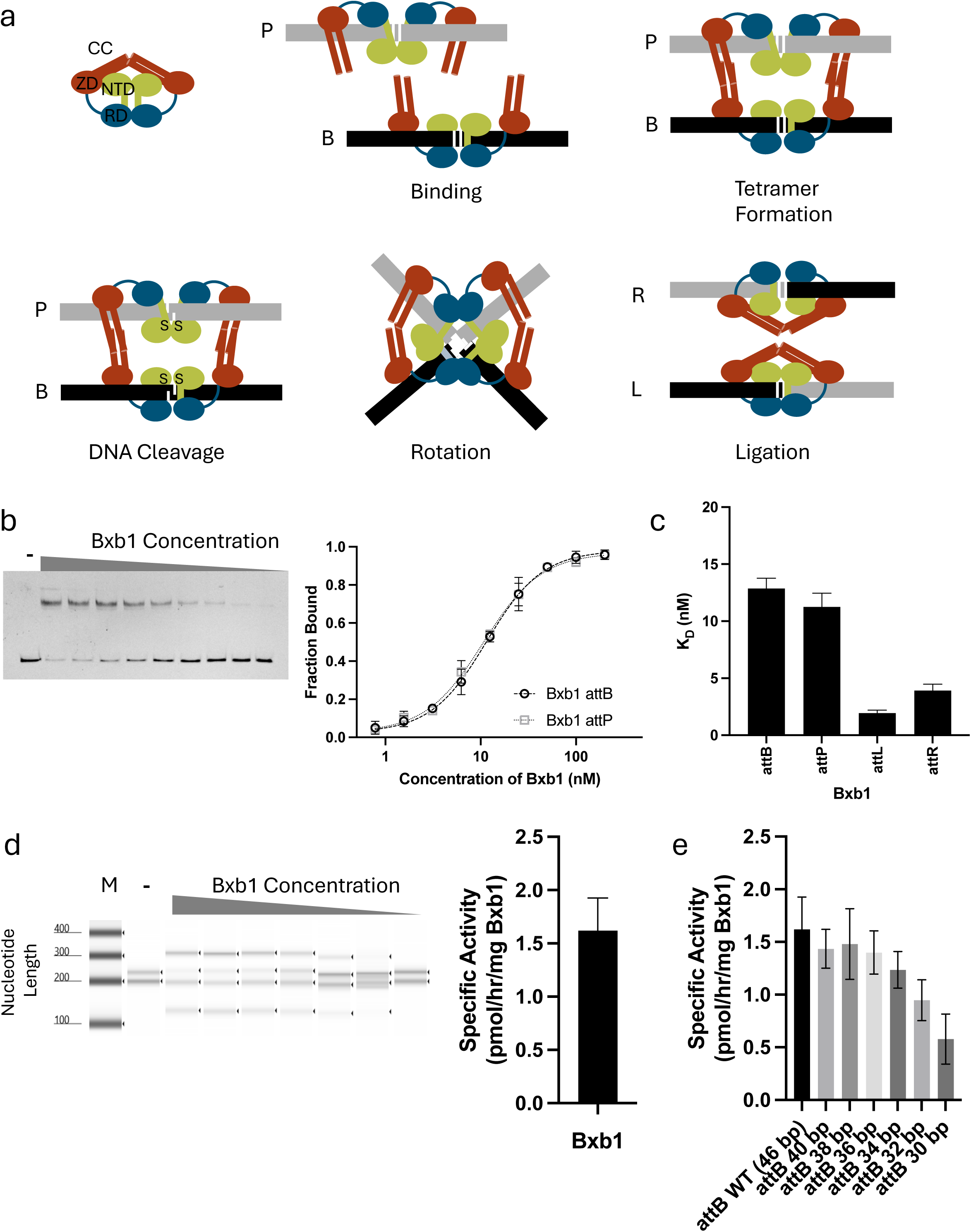
Bxb1 binding and activity assays. **a.** The Bxb1 protein has three distinct domains, a N-terminal Domain (NTD) which is responsible for both dimerization and catalysis, a Recombinase Domain (RD) and a Zinc Ribbon Domain (ZD) which contains an extended coiled-coil motif (CC). In solution Bxb1 forms a dimer which binds DNA on either side of a central dinucleotide of either and attB or attP recognition sequence. The CC domains of the attB and attP bound forms interact in the tetramer conformation where a catalytic serine residue cleaves the phosphate backbone of the DNA and forms a covalent bond. Strand exchange then occurs by rotation of the dimers around a planar interface forming two new stable-bound dimers bound to the attL and attR recognition motifs. Finally DNA ligation and dissociation leave an intact double-stranded DNA. **b.** Bxb1 DNA binding Gel shift assay with the lower band corresponding to unbound and the upper band to bound with a slow migrating species visible in the two highest concentration lanes (left). Curves showing the quantified fraction of Bxb1 bound to DNA as a function of the concentration of enzyme. n=2 error bars are standard deviation (right) **c.** Dissociation constants calculated from gel-shift binding assays with error bars representing the 95% confidence interval. **d.** Bxb1 activity assay. Activity was measured by using DNA substrates containing either attB or attP with the attachment site placed close to 3’ or 5’ end respectively. Prior to recombination the DNA fragments were 180 bp (attB) or 220 bp (attP) and upon recombination form 286 bp (attL) 115 bp (attR), size differences can be seen on the TapeStation gel image (left) and used to calculate a specific activity (right). Error bars represent the standard deviation of n=2 measurements. **e.** Specific activity measurements for Bxb1 against DNA substrates containing truncated attachment sites. For each substrate the total length of the fragment was maintained by changing the attachment sequence with A <-> C and G<->T. Error bars represent the standard deviation of n=2 measurements.

In its native environment, *Mycobacterium smegmatis,* the Mycobacteriophage protein Bxb1 catalyzes the integration of its own phage genome into the genome of the mycobacterium (17). At a biochemical level, a Bxb1 dimer sits on either of the DNA substrates (either attP or attB) and then forms a tetramer bringing the two substrates into proximity (18). A serine in the active site of Bxb1 attacks the phosphodiester backbone creating a covalent phosphoserine linkage between all strands of the DNA and rotation of the two half-sites leads to the ligation of the DNA backbones and formation of attL and attR (18,19). Using a gel-based assay and substrates of varying length we can measure the specific activity of LSI’s. Under saturating substrate conditions, Bxb1 can catalyze this reaction with a specific activity of 1.5 pmol/hr/mg Bxb1 (Fig. 1d). Further, in the absence of an accessory protein, Bxb1, like other LSIs, only catalyzes the forward reaction (20).

The DNA sequence of the substrates for Bxb1 has been well defined but the minimal attB sequence required for full recombination activity was unknown (17). We created a series of attB substrates that were all the same length (180 bp) but had the sequence of the attB mutated from either end. By measuring the activity of each of the substrates we were able to demonstrate that a 34-base-pair attB leads to a similar specific activity as the full-length, 46-base-pair attB (Fig. 1e).

### Mutations in the zinc ribbon domain affect Bxb1 activity in HEK293FT cells

The zinc ribbon domain of Bxb1 is a compact DNA binding structure that is formed from two disparate regions in the linear sequence of Bxb1, with the coiled-coil domain inserted in the middle (Fig. 2a) (10,11,21). This organization is highly conserved among LSIs. Using the published structure of the *Listeria innocua* integrase, we designed a series of point mutants within the zinc ribbon domain to probe the interaction between Bxb1 and its substrates (10,21). These mutations were focused on a series of alanine and glycine residues in a loop within the zinc ribbon domain (Fig. 2b).

**Figure 2.**
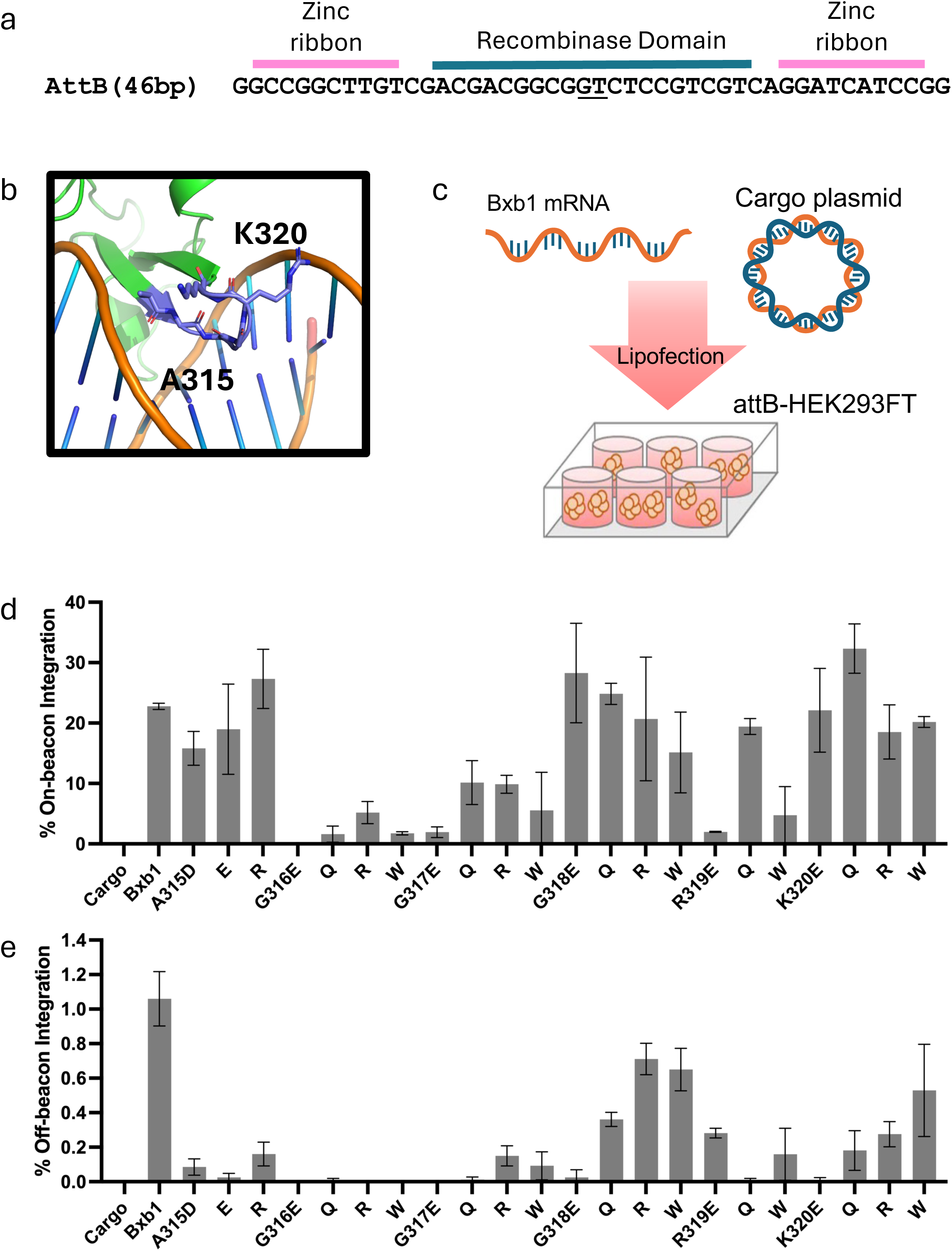
*In vitro* activity of WT Bxb1 and zinc ribbon mutants of Bxb1. **a.** The WT attachment site (attB) for Bxb1. The central dinucleotide is underlined, the recombinase binding region is indicated by a blue bar and the zinc ribbon binding region is indicated by pink bars. **b**. A close up on a mode of the zinc ribbon loop of Bxb1 that has been subject to mutation based on the structure of the LI integrase, positions A315 and K320 are indicated. **c**. Schematic of the *in vitro* experiment: Bxb1 (or Bxb1 mutants) are delivered as mRNA, insertion cargo as DNA into HEK293FT cells with a preplaced attB attachment site. **d.** On-beacon integration activity of the 22 zinc ribbon mutants as measured by ddPCR. **e.** Off-beacon integration activity of the 22 zinc ribbon mutants as measured by ddPCR at CAS031.

Mutation in the zinc ribbon domain had variable effects on integration of a DNA cargo by Bxb1. Following transfection of cargo DNA and Bxb1 mRNA into HEK293FT cells with a preplaced attB (here called ‘beacon’), we probed for on-beacon integration using ddPCR (Fig. 2c). Sub-saturating levels of both the DNA cargo and Bxb1 mRNA were used to highlight any differences amongst these Bxb1 variants. Under these conditions, we observed about 20% integration of a DNA cargo with a wild-type Bxb1 construct (Fig. 2d). We assessed the impact of mutations in the zinc ribbon domain to on-beacon integration by engineered Bxb1s using the same assay. At certain positions, like A315 and R319, we saw reduced activity for some mutants but retained (or improved) activity for other mutations (Fig. 2d). Some positions, including G316 and G317, did not seem to be amenable to mutation or at least not for the amino acids we tested. Finally, at positions G318 and K320, we found that a range of mutations was tolerated.

Comprehensive cataloguing of Bxb1 substrate preferences has identified a number of cryptic sites in the human genome that Bxb1 can use as a site for integration (15). Many of the sites results in very low levels of integration but could still be problematic when using Bxb1 in a therapeutic context (7,9,15). In HEK293, the most prominent site of mis-integration is a site on chromosome 6, CAS031 (cryptic attachment site 31) (15). We probed the extent to which zinc ribbon Bxb1 mutants can integrate at this site to identify an enzyme that has high on-beacon and reduced off-beacon integration (Fig. 2e).

Again, mutation in the zinc ribbon domain of Bxb1 had a variable effect on integration into this off-beacon location. Probing for integration at CAS031 using wild-type Bxb1 leads to about 1% integration under these experimental conditions (Fig. 2e). All the engineered Bxb1 constructs were less active on CAS031 than wild-type Bxb1, though some still retained significant activity. However, mutations at A315 appeared to significantly reduce off-beacon activity at CAS031 while retaining on-beacon integration. This is also true for G318 and K320 to glutamic acid and R319 to glutamine.

### Biochemical characterization of zinc ribbon domain mutants

We characterized a series of these mutants biochemically, measuring both binding affinities and specific activities for mutations at A315 and K320. Many mutations did not significantly impact the interaction between Bxb1 and its substrates. However, in many cases any changes we engineered led to modest decreases in the affinity between Bxb1 and attB or attP (Fig. 3a). Interestingly, the mutation of K320 to glutamic acid greatly impacted the interaction. We observed that the affinity of Bxb1 for attB increased by ∼2-fold while the affinity for attP decreased by over 20-fold. Additionally, we observed an additive effect of reduced affinity for attB and attP in the A315R and K320R double mutant.

**Figure 3.**
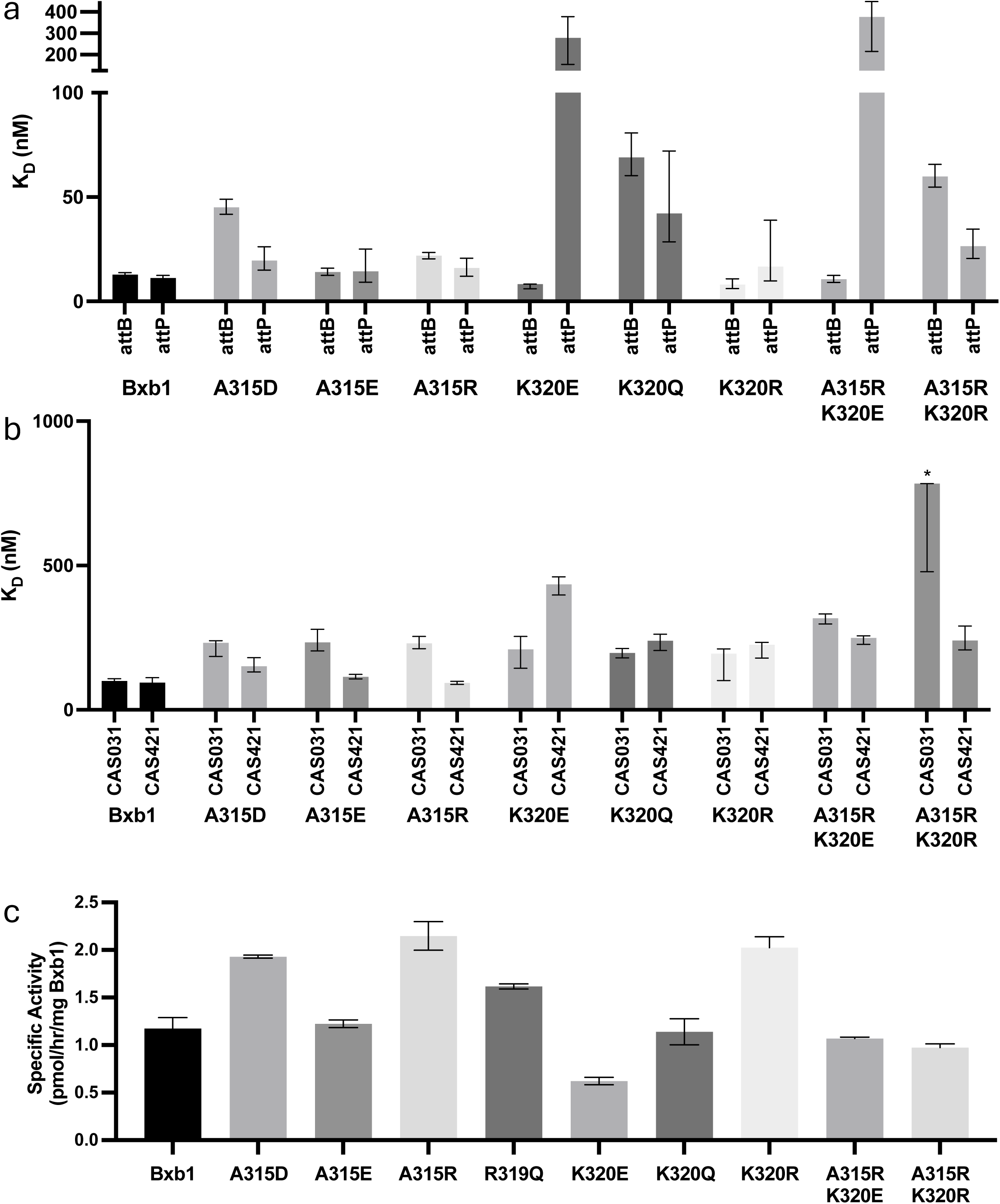
Binding and activity measurements of engineered Bxb1 mutants. **a,b.** Dissociation constants for engineered Bxb1 variants against its native substrates (a) and off-beacon substrates (b). Error bars represent the 95% confidence interval. **c.** Specific activity measurements for engineered Bxb1 variants using attB and attP containing substrates. Error bars represent the standard deviation of n=2 measurements. * Upper bounds undetermined for the A315R K320R K_D_ for CAS031.

We also investigated the impact of mutations in the zinc ribbon domain on Bxb1’s off-beacon activity. Here we looked at two different identified sites, CAS031 and CAS421, which is the most frequent off-beacon insertion site in human iPSCs (15). Using two sites allowed us to probe the effect of different mutations more thoroughly.

All tested mutations in the zinc ribbon domain led to a decrease in binding affinity (increase in K_D_) relative to wild-type Bxb1 binding to off-beacon sequences. (Fig. 3b). However, some mutations had a more significant effect on one or the other off-beacon sequences. For instance, K320E led to a 2.1-fold weaker K_D_ for CAS031 compared to wild-type Bxb1 and a 4.6-fold weaker K_D_ for CAS421. Conversely, A315R had minimal effect on binding to CAS421 compared to wild-type Bxb1 but led to a 2.3-fold weaker K_D_ for CAS031. The intermediate effect on binding affinity observed in the A315R/K320E double mutant suggests that DNA recognition by Bxb1 is more complex than can be predicted by simple mutations.

We measured the specific activity for the same series of mutants but did not see any significant differences for many of the constructs (Fig. 3c). Even for K320E, a modest reduction in the specific activity was observed. However, it is likely that under conditions of limiting substrates a more pronounced effect on activity may be seen for K320E. While A315R or K320R mutations led to an increase in activity, no additive effect was observed in the double mutant but rather a decrease in specific activity was detected. Overall, the A315R single mutation led to the greatest increase in activity.

### Naturally occurring variants of Bxb1 are as active as wild-type Bxb1 on Bxb1 substrates

There are almost 200 sequences deposited in the NCBI database that share 90% sequence similarity to Bxb1. These sequences deviate from Bxb1 at positions through the protein sequence (Fig. 4a). However, they are enriched within the DNA binding domains of Bxb1. In fact, very few positions in the NTD or coiled-coil seem to differ from the canonical Bxb1 sequence. While we have not mapped the native attB or attP sequences for these new LSIs, we decided to assess whether any of these had activity on the original Bxb1 substrates.

**Figure 4.**
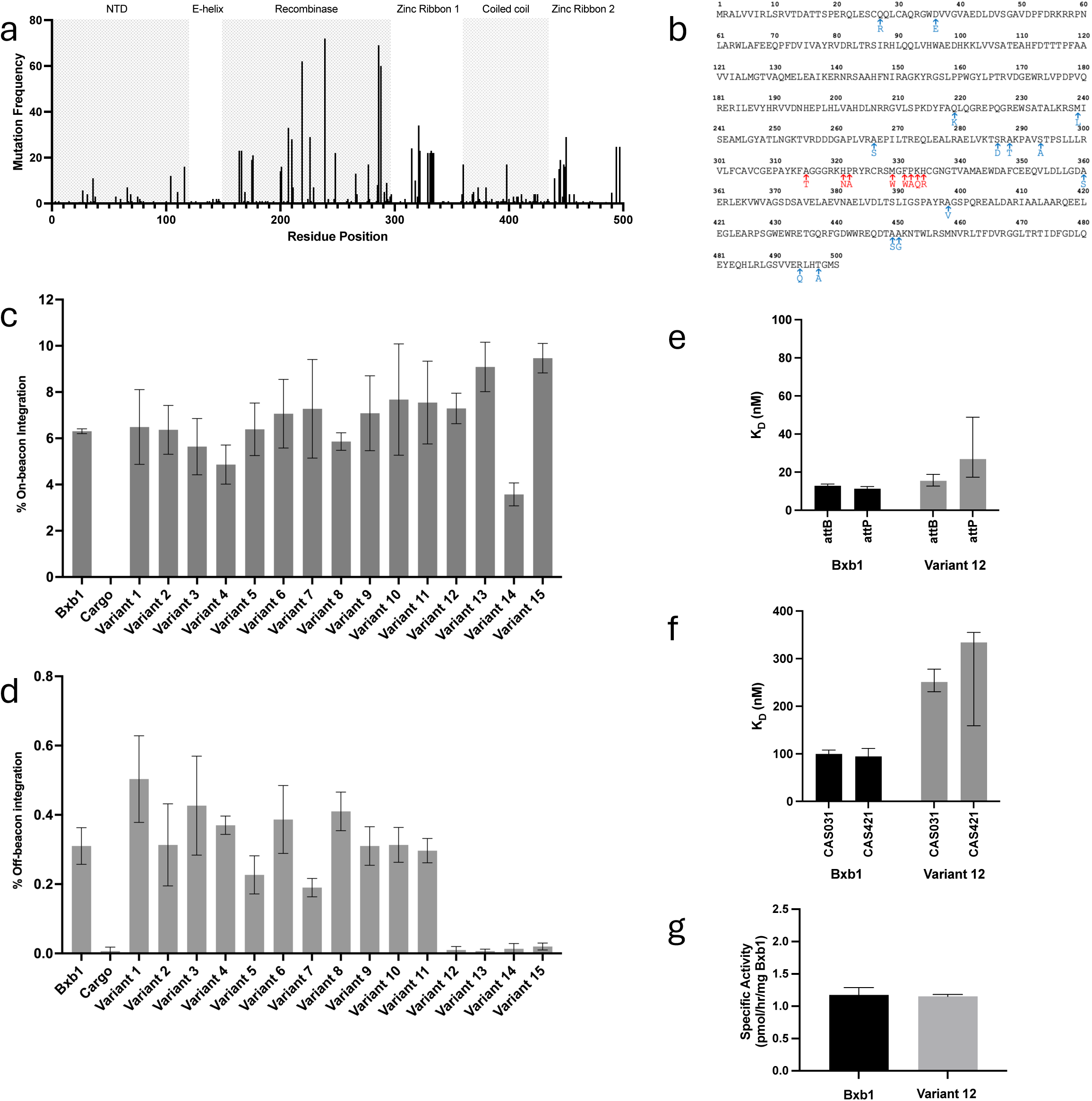
Activity of variants of Bxb1. **a.** Table of the frequency of a mutation at each amino acid position within Bxb1. **b.** Location of the specific mutations within the 15 tested naturally occurring variants indicated by arrows and amino acid changes on the WT sequence of Bxb1. The red arrows highlight the constellation of mutations in the zinc ribbon domain that were used to generate variants 12, 13, 14, and 15. **c.** On-beacon integration activity of the 15 naturally occurring mutants as measured by ddPCR. Error bars represent the standard deviation of n=3 measurements. **d.** Off-beacon integration activity of the 15 naturally occurring mutants as measured by ddPCR at CAS031. Error bars represent the standard deviation of n=3 measurements. **e,f.** Dissociation constants for Bxb1 variant 12 against attB and attP (e) and off-beacon substrates (f). Error bars represent the 95% confidence interval, n=2. **g.** Specific activity measurements for engineered Bxb1 variant 12 using attB and attP containing substrates. Error bars represent the standard deviation of n=2 measurements.

Using a HEK293FT cell line with a pre-installed beacon, we assessed the activity of fifteen naturally occurring Bxb1 variants. These variant proteins had between one and nine differences from wild-type Bxb1, distributed throughout the protein sequence (Fig. 4b). Interestingly, most of the naturally occurring variants, unlike our designed variants, were at least as active, if not more active, than wild-type Bxb1 (Fig. 4c). We observed the biggest differences with mutations clustered in the zinc ribbon domain.

We also probed these naturally occurring variants for their off-beacon integration activity at CAS031. Unlike the engineered Bxb1 enzymes, most of these variants retained significant off-beacon activity (Fig. 4d). However, variants 12, 13 and 15 showed on-beacon activity comparable to wild-type Bxb1 and exhibited low or no activity off-beacon at CAS031. All of these variants contain the same constellation of mutations in the zinc ribbon domain (A315T, H321N, P322A, M329W, F331W, P332A, K333Q, H334R). Variant 13 also contains S293A and variant 15 contains A360S.

We measured the binding affinity and specific activity for variant 12. We saw very subtle differences in binding affinity for wild-type attB or attP (Fig. 4e). We also measured binding of variant 12 to both CAS031 and CAS421, where we saw a significantly weaker binding affinity to both sequences compared to wild-type Bxb1 (Fig. 4f). We did not see any difference in the specific activity between variant 12 and wild-type Bxb1 (Fig. 4g). However, many of the altered positions between the two proteins would likely mediate interactions between the DNA and the LSI. Additionally, the natural substrates for these variant Bxb1 proteins may be different from those of canonical Bxb1.

### Engineered Bxb1 variants improve Bxb1 activity *in vivo*

For genome engineering efforts, LSIs must be active *in vivo*. Using transgenic mice that had been generated to carry an attB in their genome, we assessed the activity of engineered Bxb1 variants on hepatocytes *in vivo* (Fig. 5a). Delivery was facilitated by an IV injection of an LNP carrying the mRNA encoding the LSI and template DNA was delivered by a self-complementary AAV8 contain the coding sequence for the human F9 gene. Our early studies found that Bxb1 showed only modest activity *in vivo,* with wild-type Bxb1 (containing a nuclear localization signal (NLS) and an HA-tag or just an NLS) resulting in only 1-2% integration (Fig. 5b, Bxb1NLS-1, Bxb1 NLS-2). We decided to apply traditional protein engineering efforts to improve this, namely by potentially increasing the stability of the protein through addition of stability tags. An N-terminal fusion with a STABILON tag (derived from the p54/Rpn10 ubiquitin receptor subunit of the *Drosophila* proteosome) (22) (Fig. 5b, Bxb1 Stabilized-1) increased in vivo potency almost 3-fold. Combining the STABILON tag with a second published sequence, Exin21 (Fig. 5b, Bxb1 Stabilized-2) further increased in vivo potency (23). Combining these two tags on the N-terminus of Bxb1 NLS-1 lead to over 5% gene insertion.

**Figure 5.**
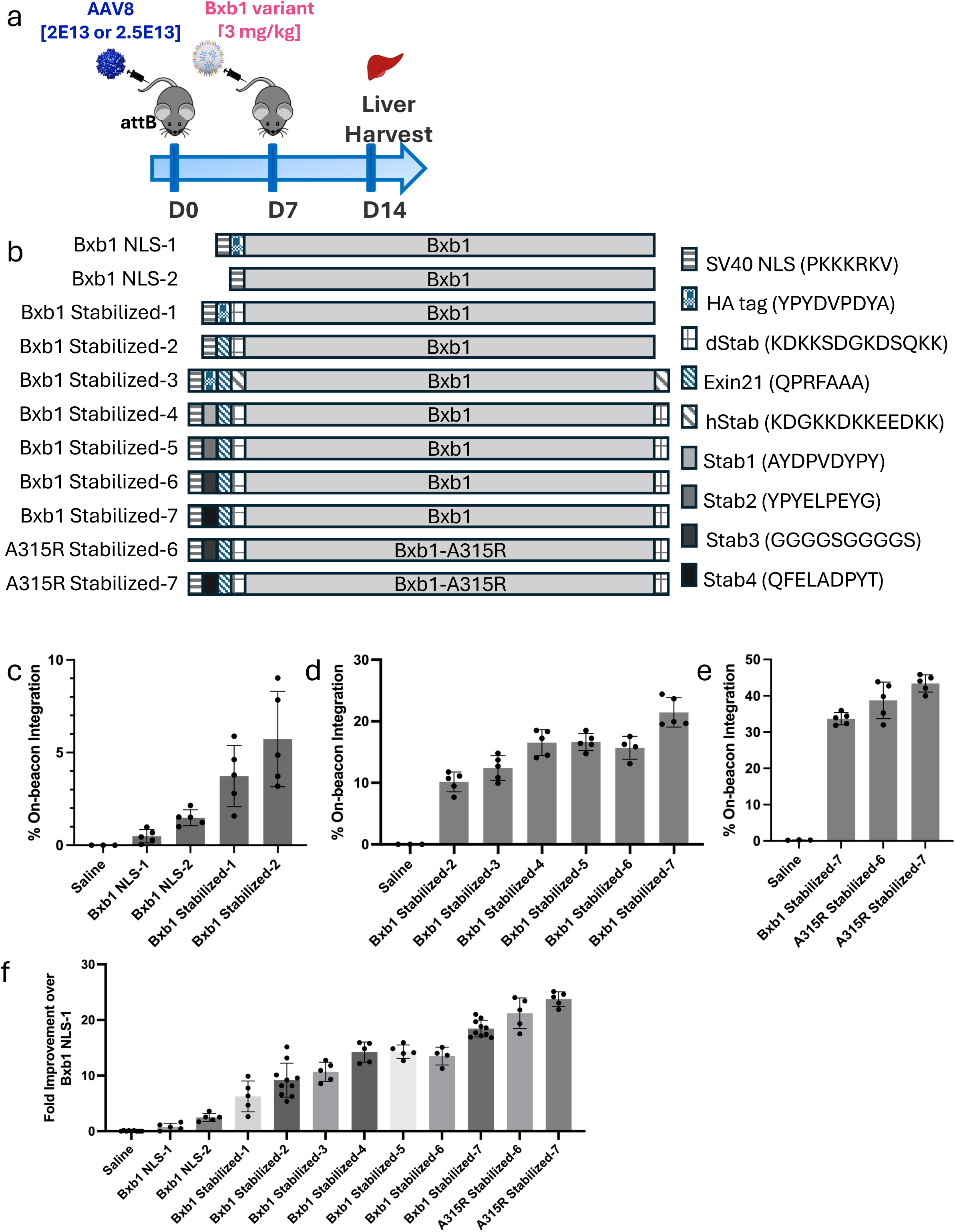
In vivo activity of stabilized Bxb1. **a.** Schematic of the *in vivo* experiment: On day 0, AVV containing a DNA cargo is delivered via IV. On day 7, Bxb1 (or stabilized Bxb1) is delivered via LNP. On day 14, livers are harvested and genomic DNA is probed for on-beacon integration. **b.** Schematic for the amino acid sequences of the stabilized Bxb1 molecules. Different stability tags are indicated by variable shadings. **c**. *In vivo* activity for original stabilized Bxb1 constructs. **d.** *In vivo* activity for the final panel of stabilized Bxb1 constructs. **e**. *In vivo* activity for Bxb1 constructs containing both A315R and stabilization tags. **f**. Fold increase of *in vivo* activity of Bxb1 constructs compared to Bxb1 WT-1.

Following some other optimizations of components, we were able to achieve 10% insertion with Bxb1 Stabilized-2 (Fig. 5c). Addition of a second STABILON sequence to the C-terminus (Bxb1 Stabilized-3) had a modest effect on the activity of Bxb1 *in vivo* (Fig. 5d). However, we had also modified the identity of the STABILON tag to the sequence from the human proteasome subunit, which may have led to lower activity overall (22). Returning to the Drosophila STABILON sequence, we altered the sequence N-terminal of the Exin21 tag to remove the HA-tag. Bxb1 Stabilized-4-7 variants all have a series amino acids containing charged or polar residues to act as an alternative to the HA-tag. For most of these we detected an improvement over the original stabilized sequence (Stabilized Bxb1-2) although these proteins also contain an additional STABILON at the C-terminus. For Bxb1 Stabilized-7, the rate of cargo integration is almost double compared to Bxb1 Stabilized-2.

Finally, we combined our best stabilized versions of Bxb1 with our engineered Bxb1 variants. The most active engineered Bxb1 was A315R and we combined this with two of the stabilized variants of Bxb1. We were able to achieve over 40% cargo integration in bulk liver tissue using Stabilized-A315R-7 (Fig. 5e). From Bxb1 NLS-1 to our final construct, we were able to improve *in vivo* potency of the integrase by over 25-fold (Fig. 5f).

## Discussion

The ability to specifically place a large piece of DNA into a specific location has the potential to revolutionize genetic medicines. It can enable advanced cell engineering, as well as endogenous gene replacement for patients suffering from genetic diseases. Programmable Genomic Integration (PGI) based on LSIs has the potential to efficiently overcome the limitations of previous gene insertion technologies, however LSIs found in nature have challenges associated with potency and specificity. Here we described the development of a suite of novel integrases that could begin to address some of the fundamental issues with the current editing technologies.

One of the concerns with integrases is a lack of specificity (15). If gene insertion were to occur within a deleterious genomic site (e.g. tumor suppressor), it could lead to an unfavorable safety profile. The tumor suppressor gene would likely now be inactive and could result in development of malignancies. Our group and others have validated at least 50 sites of off-beacon integration by wild-type Bxb1 in mammalian cells, with many more potential sites showing detectable activity in cell-free biochemical assays (7,9,15).

To address this specificity issue, we have explored two different approaches. First, we used structure-based design to make specific point mutations in the zinc ribbon of Bxb1. Because Bxb1 seems to have fewer sequence-specific contacts in this part of the protein we hypothesized that we would be able to introduce mutations that would increase the stringency of binding to the substrates. We found several positions (notably A315 and K320) where mutation had a small or no effect on Bxb1 on-target activity in HEK293FT cells – leading to similar integration at on-beacon sites and decreased integration at off-beacon sites compared to wild-type Bxb1. These mutants positively impacted DNA binding by Bxb1, leading to proteins with higher affinity for the wild type sequence and lower affinity for the off-target sequences.

We also analyzed deposited Bxb1-like sequences to find common mutations that may be beneficial to enzyme activity due to their enrichment in these orthologues. We found one constellation of mutations that appeared to be quite beneficial for Bxb1 activity – increasing the level of on-beacon integration in HEK293FT cells while decreasing the level of off-beacon integration to virtually undetectable levels.

Large serine integrases also suffered from low levels of activity in vivo. Other groups have increased Bxb1’s activity using protein evolution schemes to introduce mutations throughout the sequence to achieve efficient integration (7,13). We have used a different method, notably protein tagging with stabilizing sequences. This has led to significant improvements in Bxb1 activity – from less than 2% integration for unstabilized Bxb1 to over 40% in vivo integration for the fully stabilized constructs. This level of integration is now sufficient to be considered for use in therapeutic genome engineering using LSIs.

Bxb1 has historically been the integrase of choice for genomic engineering efforts (12). Here we have described improvements to Bxb1 to increase specificity and activity that will enable it to continue to lead the field for large serine integrases.

## Figure Legends

**Supplemental Figure 1.**
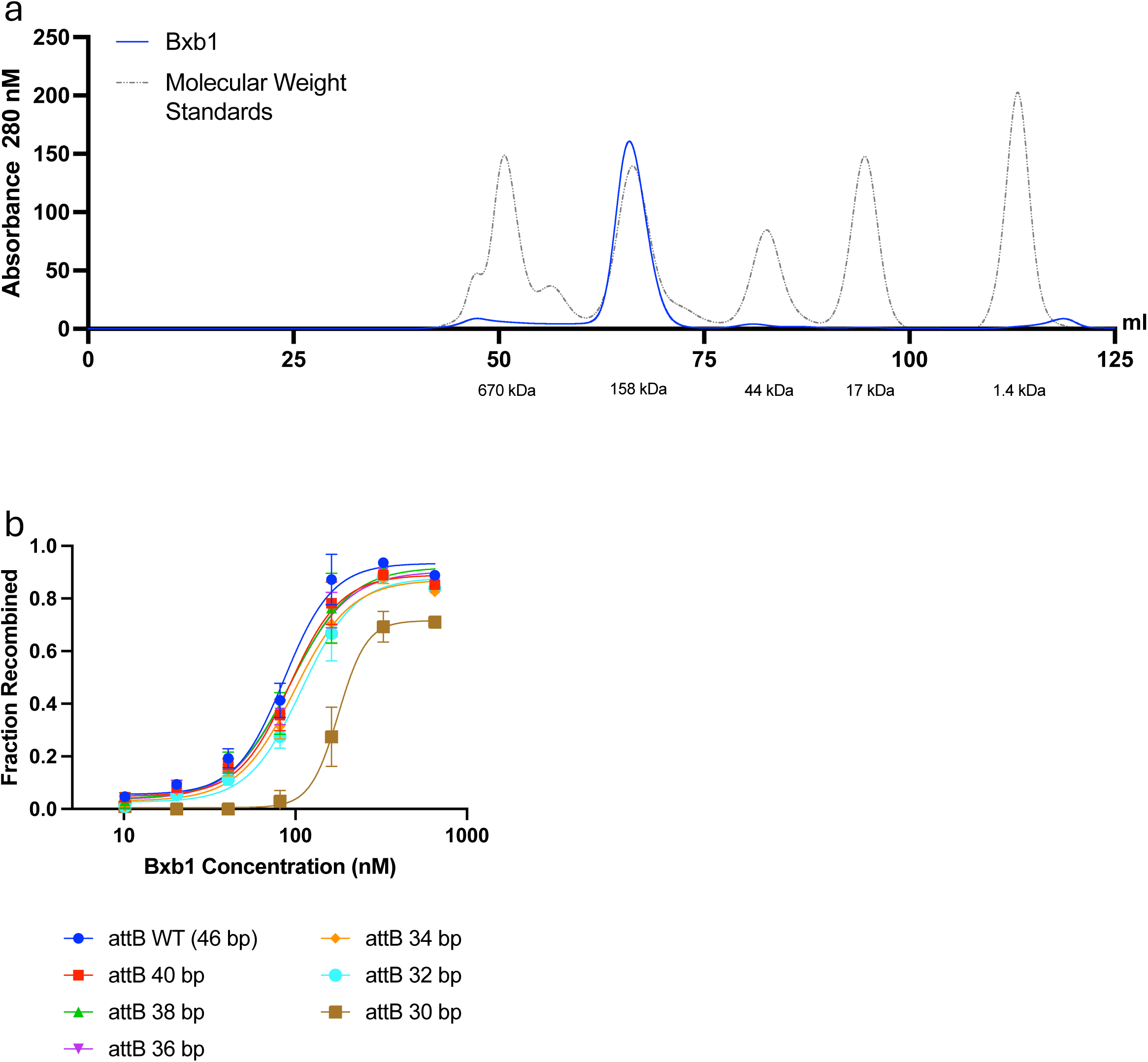
**a.** Size exclusion chromatography trace of Bxb1 2-500 showing its observed retention time co-elutes with a 158 kDa molecular weight standard. **b.** Recombinase activity assay curves for minimal attB sequence determination with the WT attP sequence. Error bars represent standard deviation n=2.

**Supplemental Figure 2.**
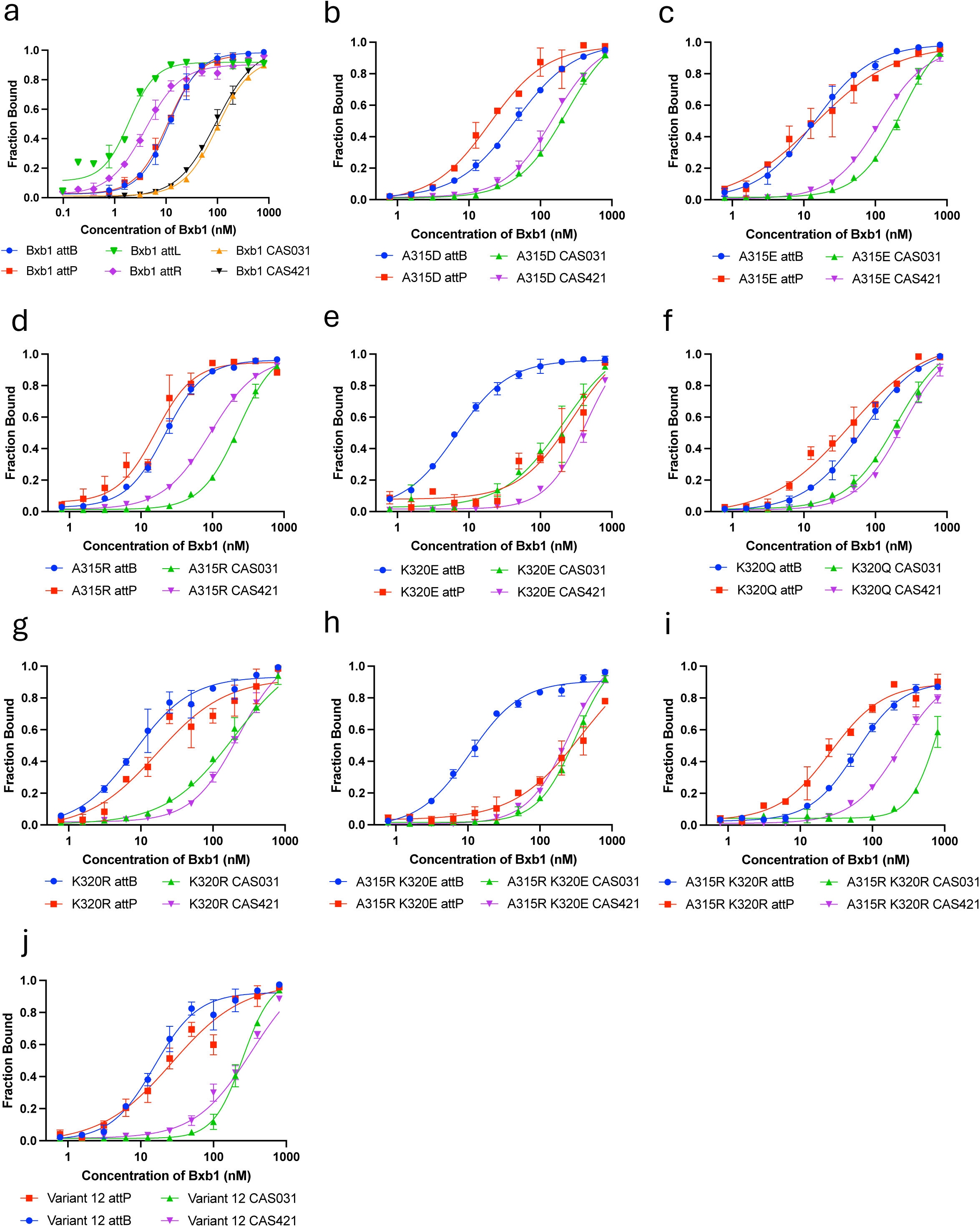
**a.** Binding assay curves for Bxb1, for attB, attP, attL, attR, and CAS031 and CAS421 off-beacon sequences. **b-j.** Binding assay curves for Bxb1 variants for attB, attP, and CAS031 and CAS421 off-beacon sequences. Error bars represent standard deviation n=2.

**Supplemental Figure 3.**
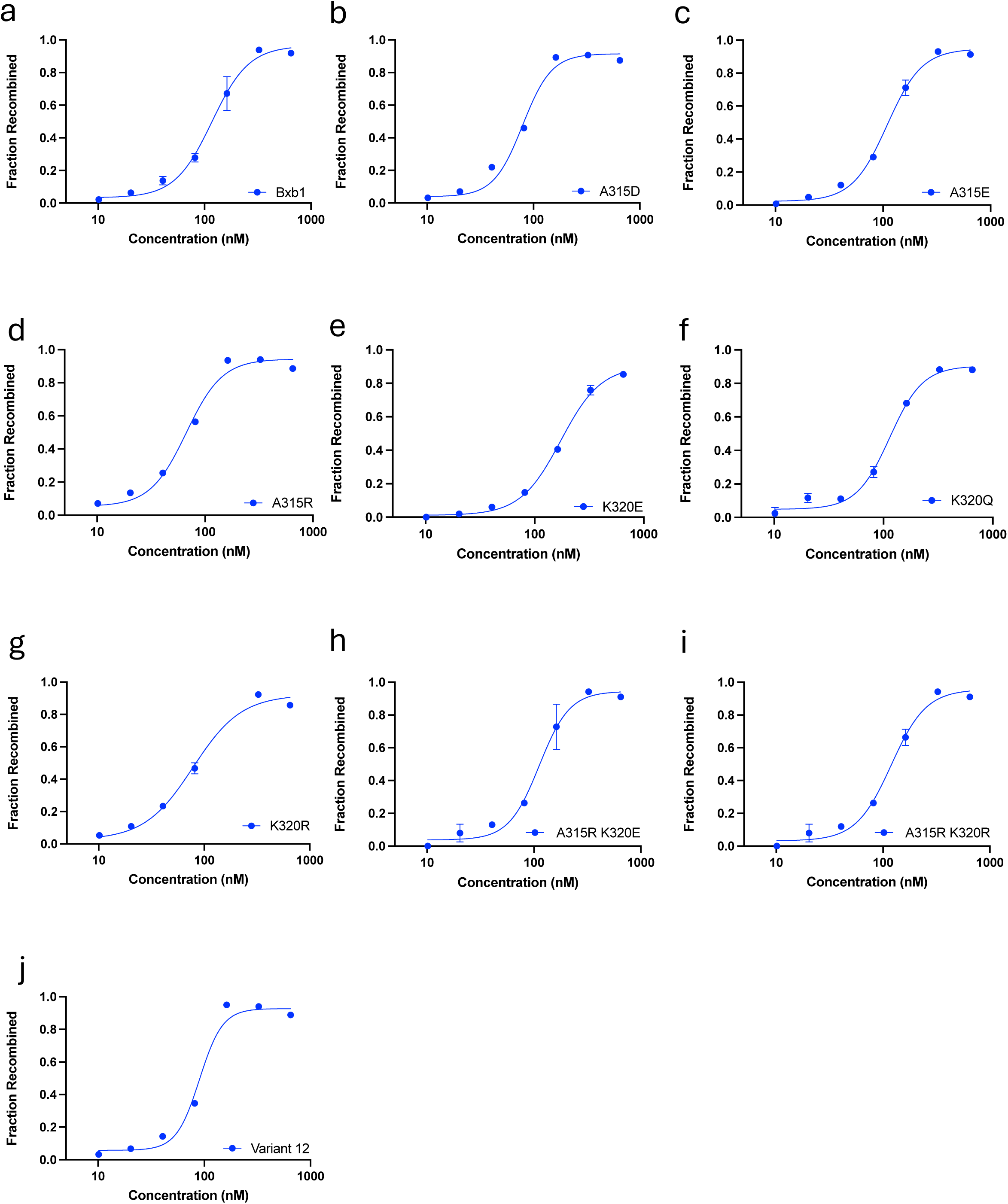
**a-j.** Recombinase activity assay curves for attB and attP for Bxb1 and variants. Error bars represent standard deviation n=2.

**Table 1.**
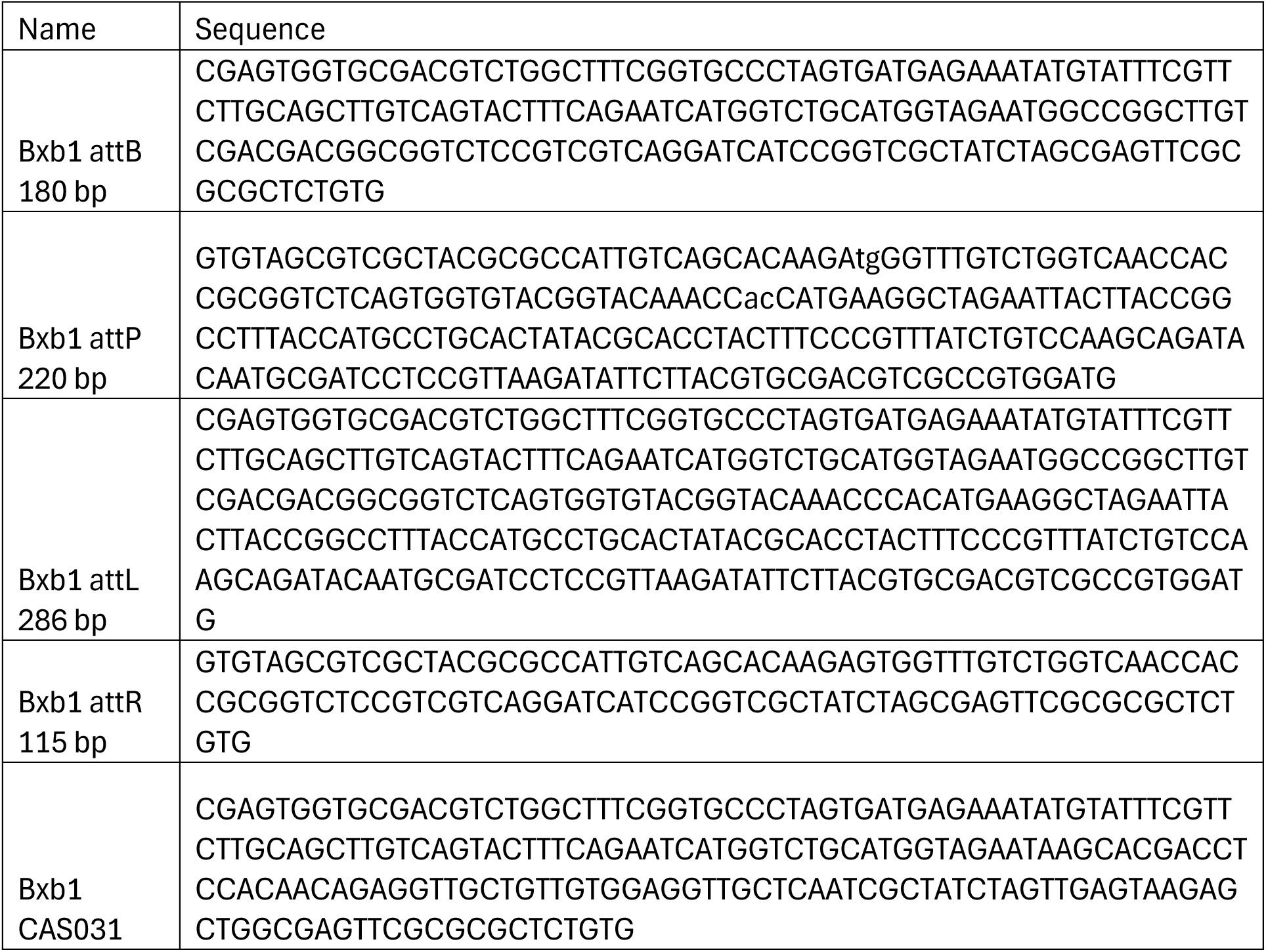
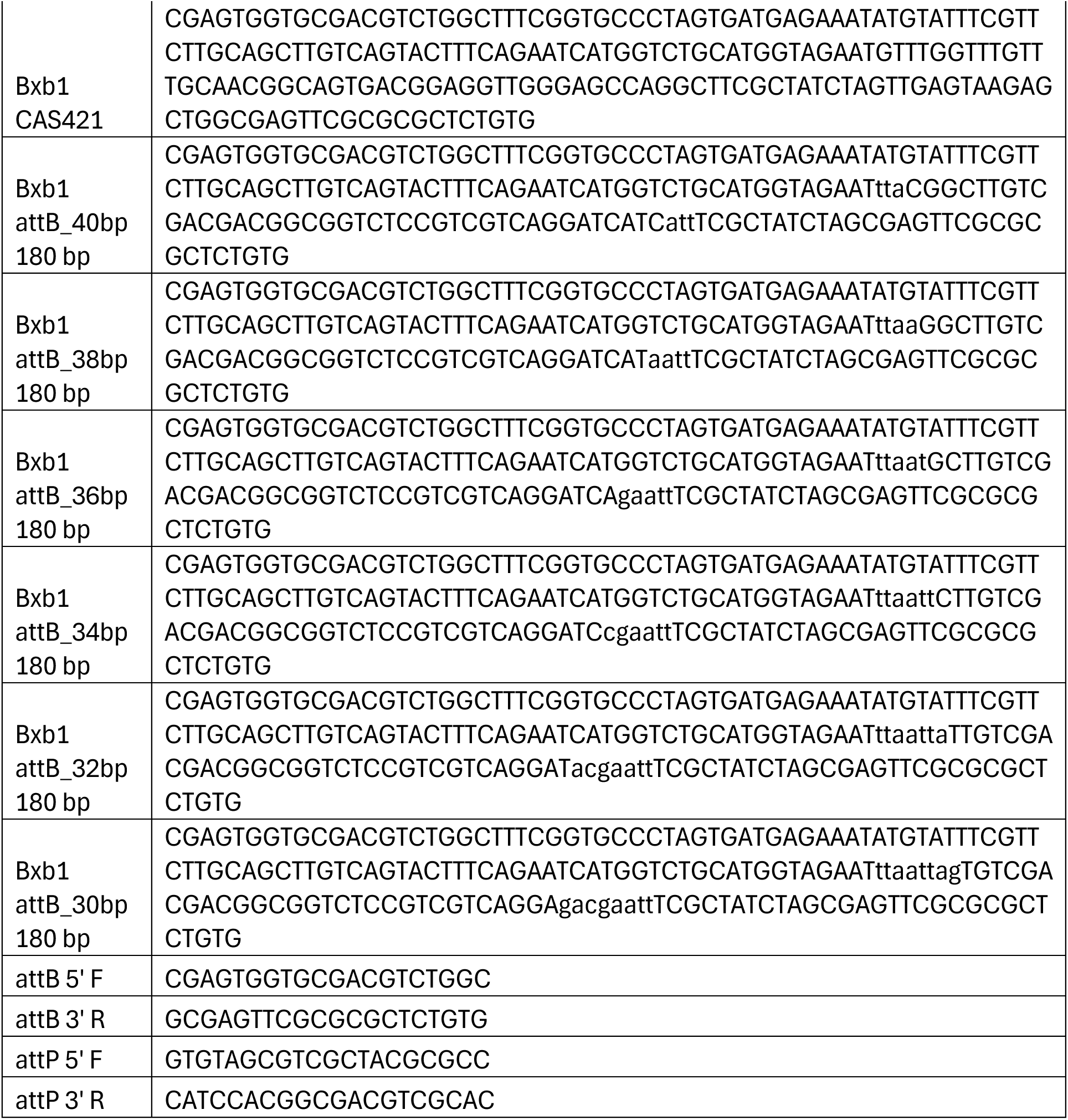
related to Figures 1b, 1c, 1d, 1e, 3a, 3b, 3c, 4e, 4f, 4g. DNA substrates for biochemical assays.

**Table 2.**
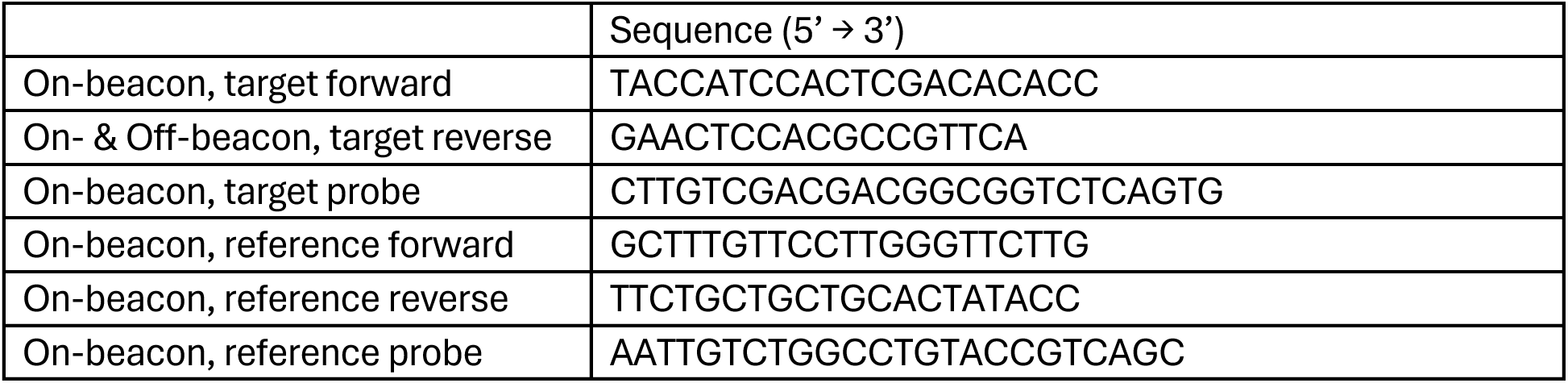
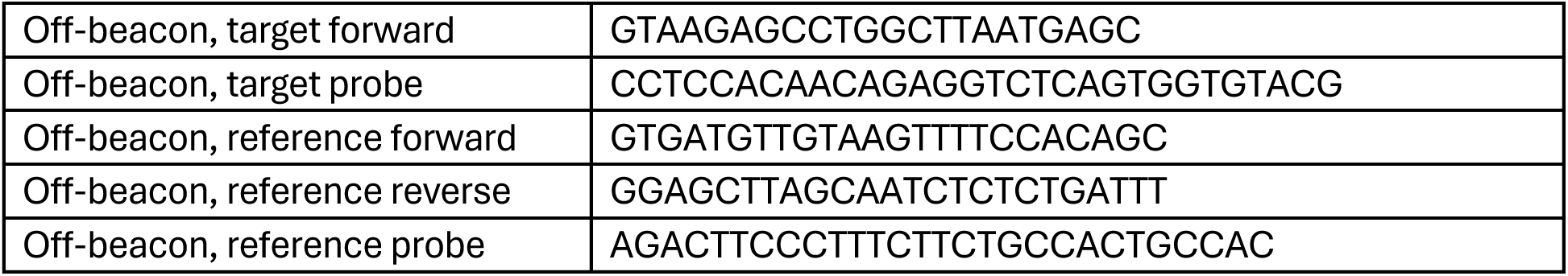
related to Figures 2c, 2d, 4c and 4d. DNA primers and probes for *in vitro* ddPCR assay.

**Table 3.**
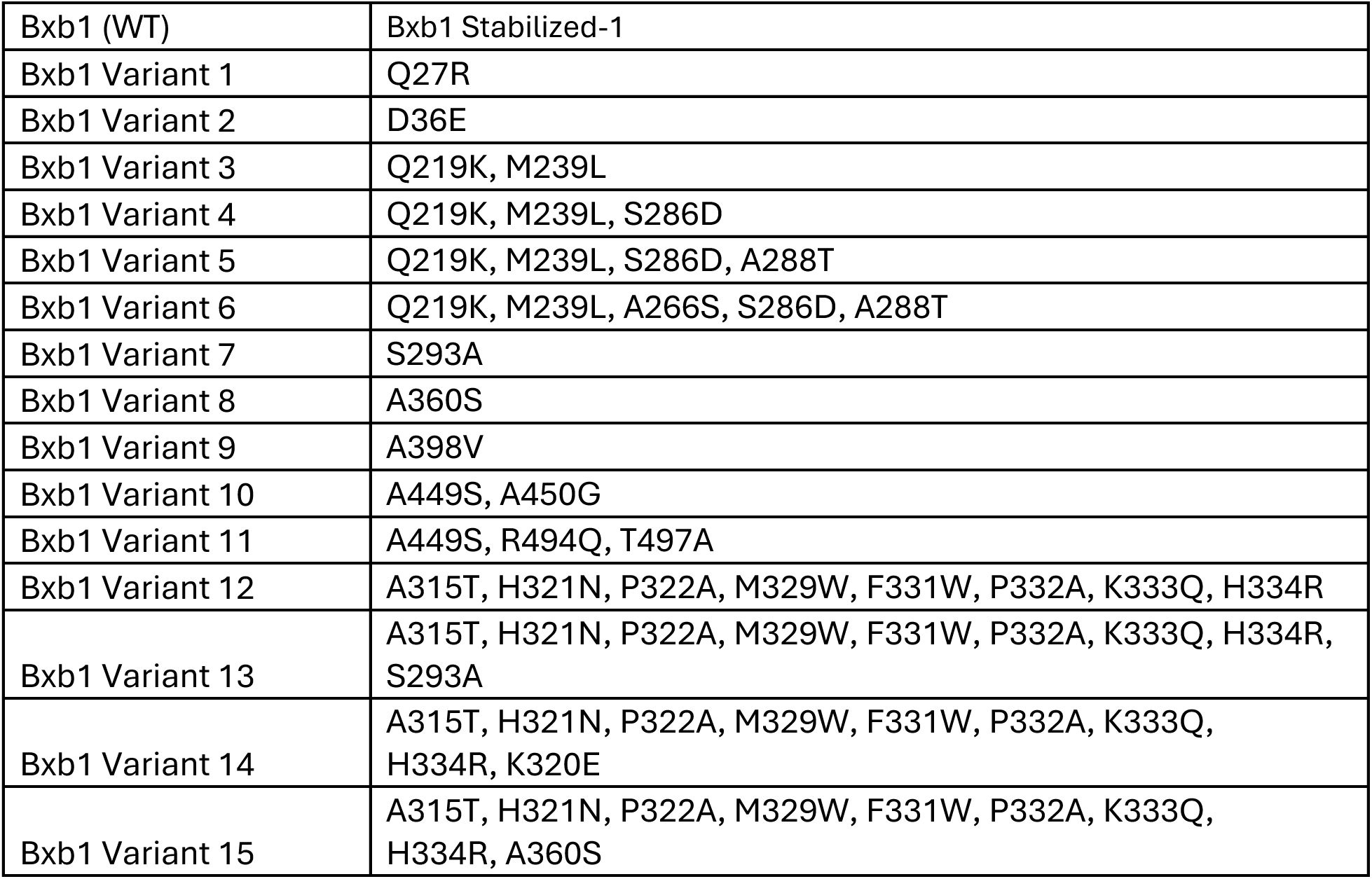
related to Figure 4c. Protein mutations of studied Bxb1 variants.

**Table 4.**
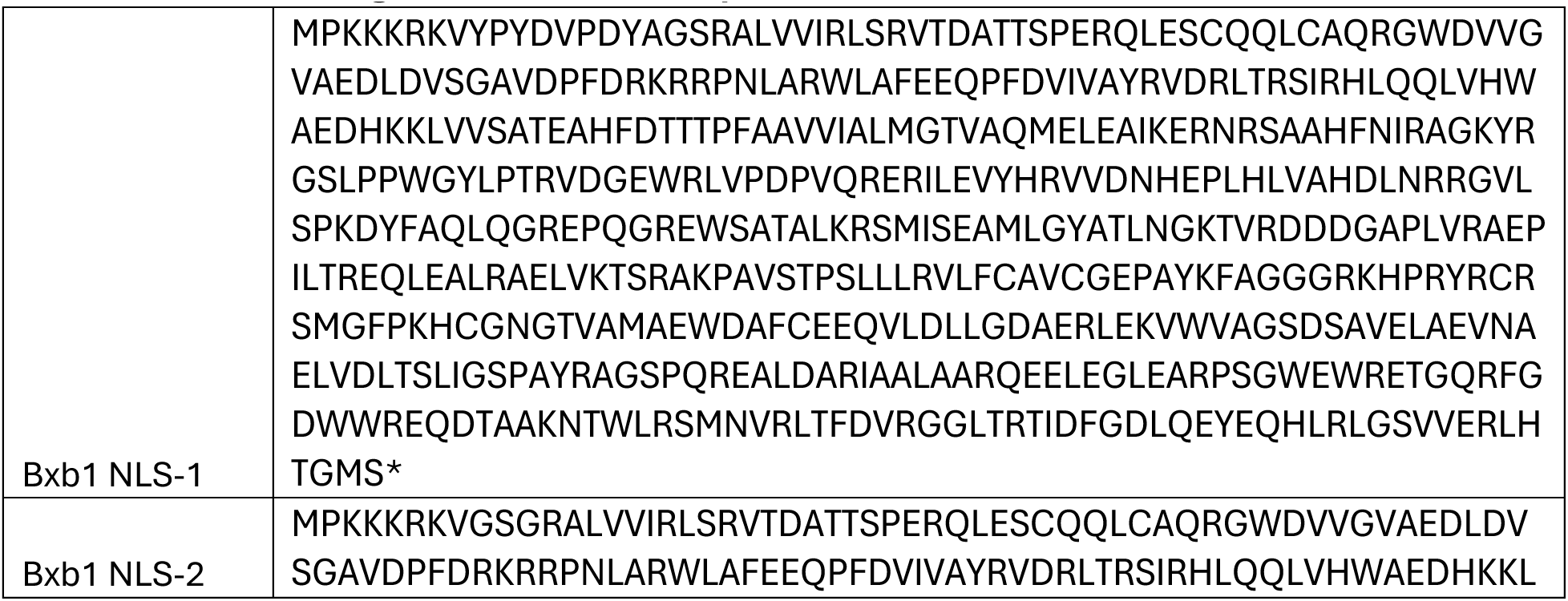
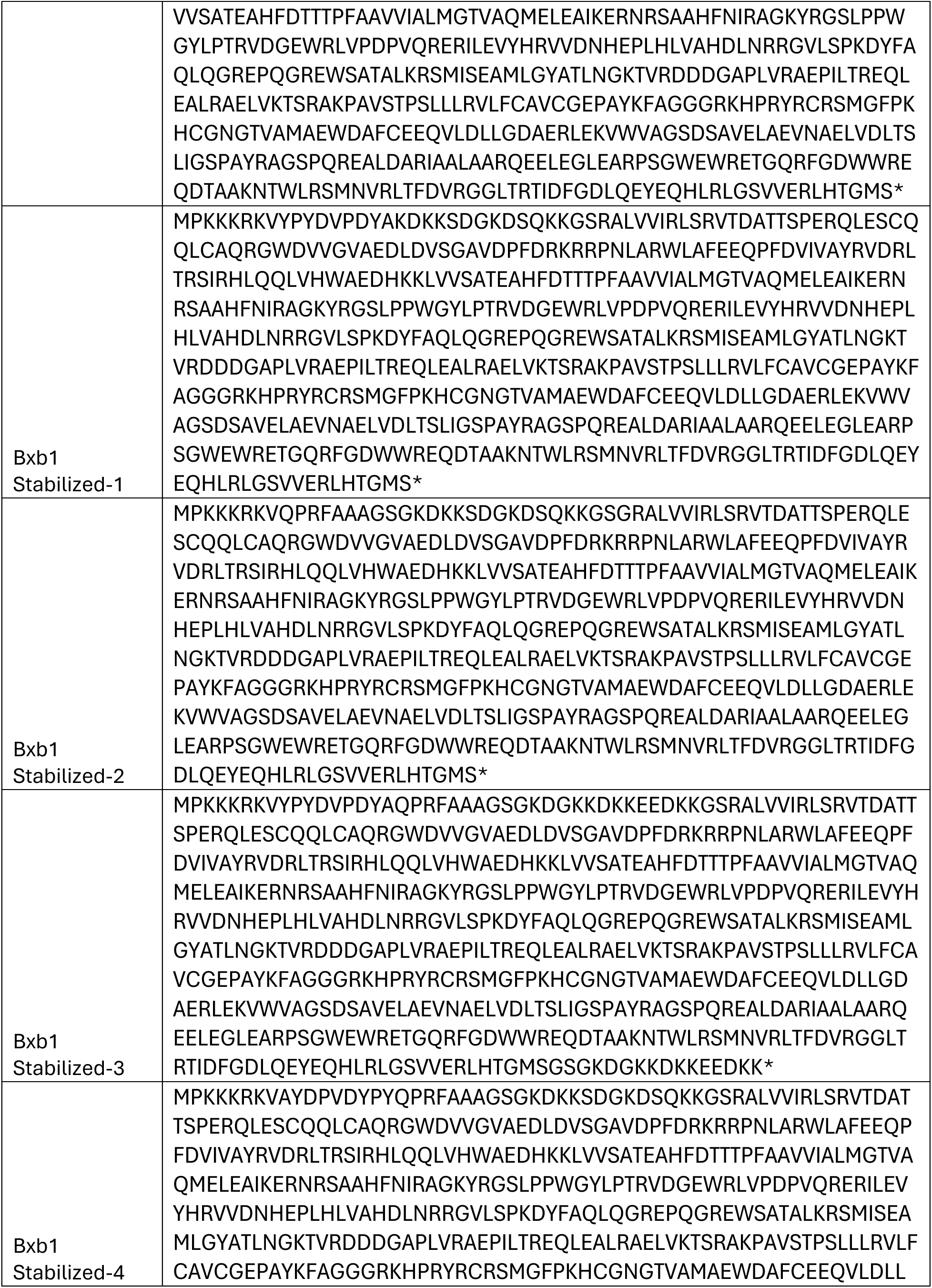
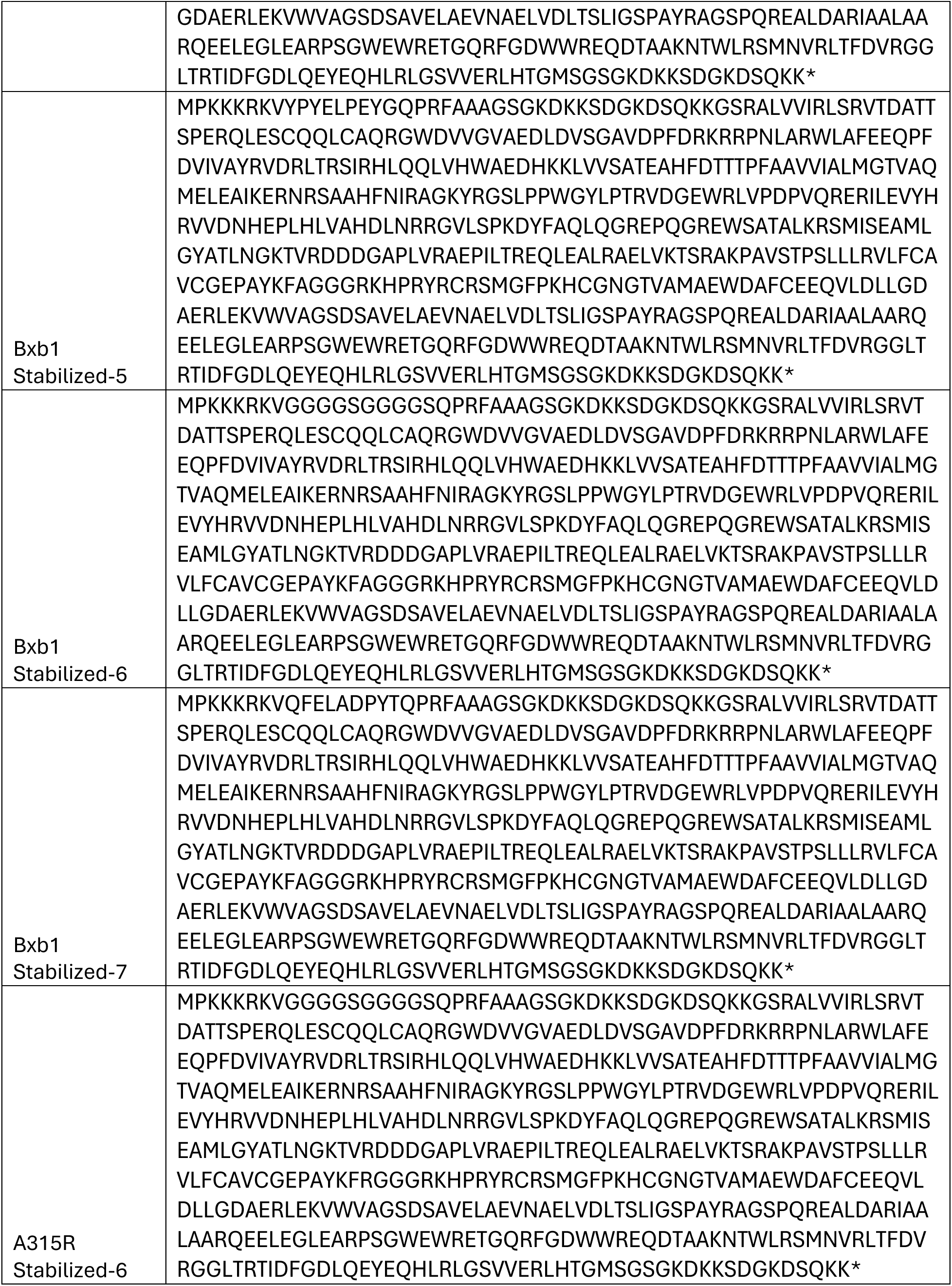
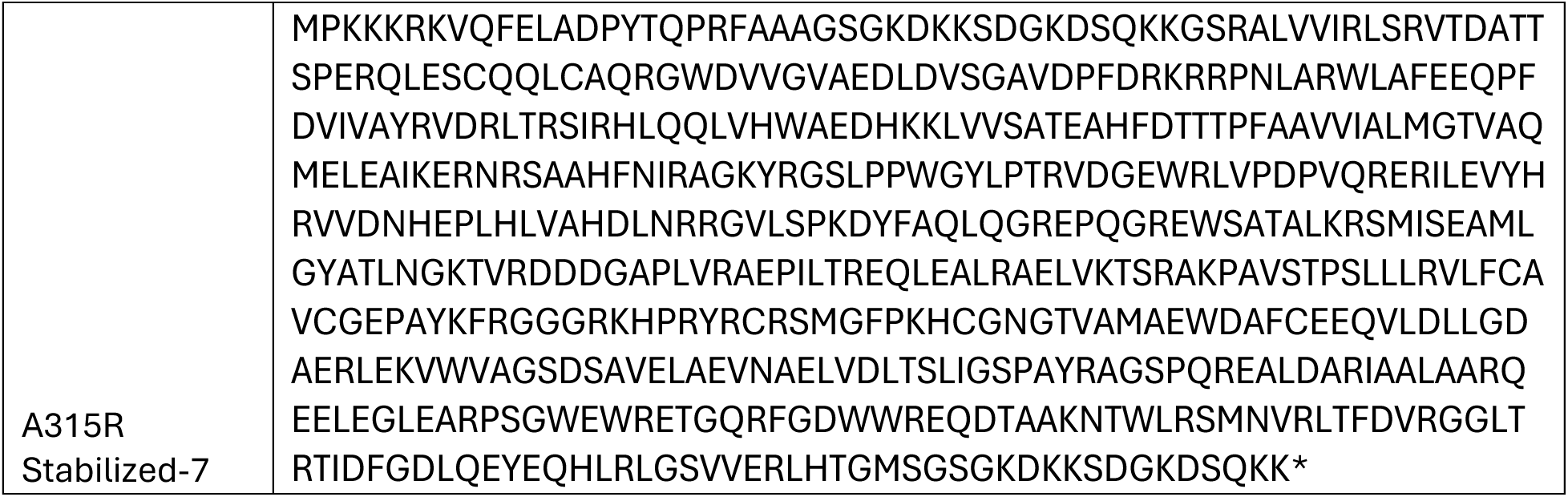
related to Figure 5b. Protein sequences for stabilized variants.

**Table 5.**
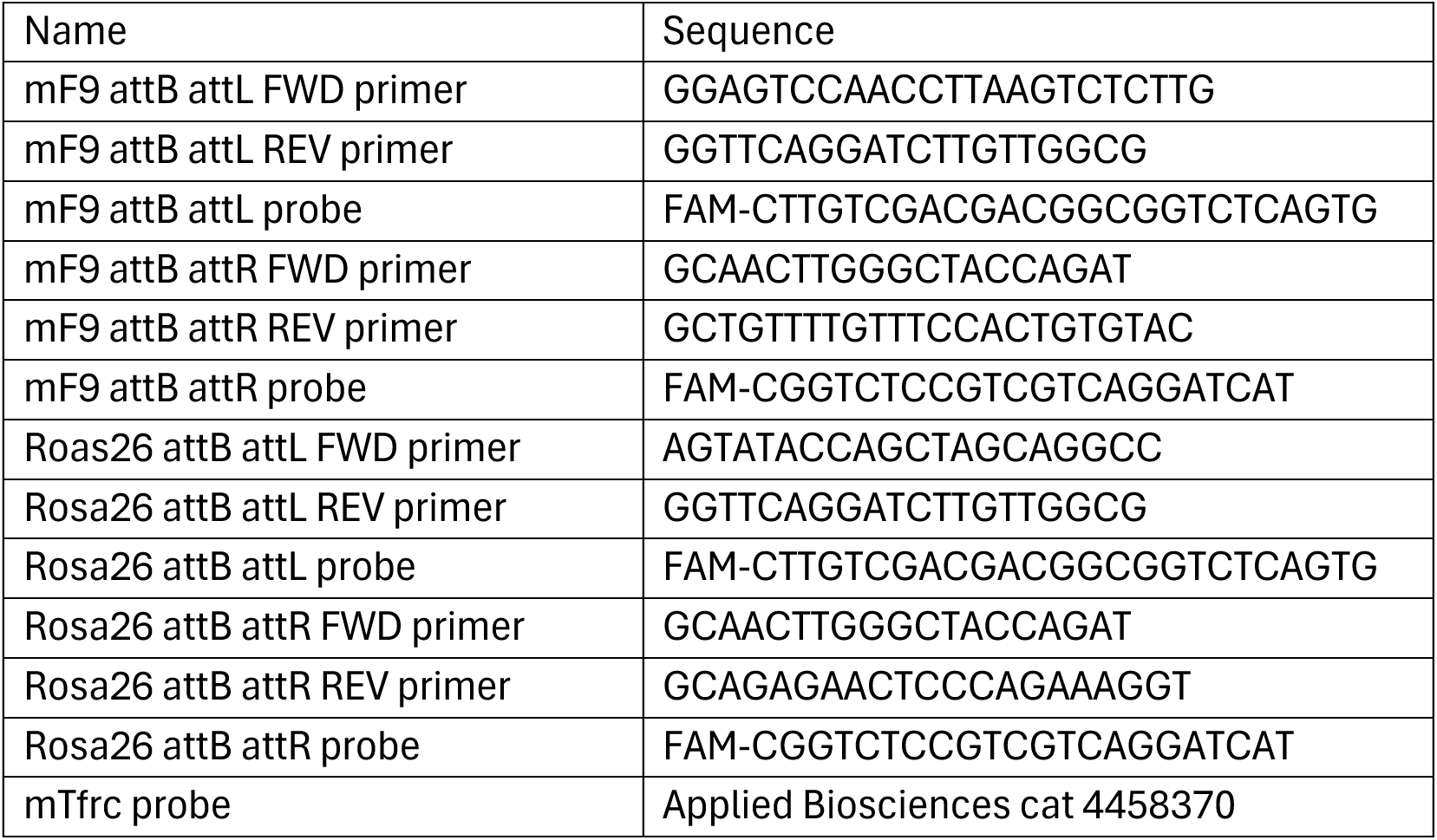
related to Figure 5c, 5d, 5e. DNA primers and probes for *in vivo* ddPCR assay.

## Method Details

### Protein Purification

Bxb1 residues 2-500 and all mutants were cloned into a pET28 expression vector containing a Thioredoxin tag, CL7 affinity tag, 8x His affinity tag, and SUMO tag (Trialtus Biosciences 20-1022). Protein was expressed in BL21(DE3) E. coli cells (New England Biolabs) grown in Terrific Broth (Teknova T7660) and induced with 1 mM IPTG for 16 hours at 18 °C. The resulting cell pellet was resuspended in Ni NTA Buffer A with Halt Protease Inhibitor (Thermo Scientific 78439) and lysed by sonication. Crude lysate was centrifuged at 30,000 x g for 30 min and the resulting cleared lysate applied to a 5 mL HisTrap HP column (Cytiva 17-5255-01). The column was washed with Ni NTA Buffer A: 50 mM Tris pH 8.0, 500 mM Sodium Chloride, 25 mM Imidazole, 1 mM DTT and eluted by a gradient with Ni NTA Buffer B: 500 mM Sodium Chloride, 250 mM Imidazole, 1 mM DTT pH 8.0. Following elution Arginine was added to reach a concentration of 200 mM and the protein was cleaved with 50 units of SUMO Protease (Trialtus Biosciences 30-1130) for 16 hours at 4°C and exchanged into Heparin Buffer A with a HiPrep 26/10 Desalting column (Cytiva 17-5087-01). Cleaved enzyme was bound to a 5 mL HiTrap Heparin HP column (Cytiva 17-0407-03) and washed with Heparin Buffer A: 50 mM Tris pH 8.0, 100 mM Sodium Chloride, 200 mM Arginine, 1 mM TCEP and eluted by gradient elution with Heparin Buffer B: 50 mM Tris pH 8.0, 1 M Sodium Chloride, 200 mM Arginine, 1 mM TCEP. Final polishing was performed using a HiLoad Superdex 200 16/600 column (Cytiva 28-9893-36) and the buffer 50mM Tris, 500mM Sodium Chloride, 200mM Arginine, 1mM TCEP.

### Biochemical Measurement of Bxb1

Substrates for biochemical assays were generated using single stranded gBlocks (IDT) designed for each specific target. Gel shift binding assays used a Cy3 or Cy5 fluorophore added to the 3’ end using a fluorescent tagged primer, biochemical activity assays used the same substrates with no fluorophore added.

Gel shift binding experiments were carried out with 1 nM substrate for K_D_ values above 10 nM and 0.5 nM substrate for K_D_ values below 10 nM. Gel shift binding buffer consisted of 20 mM Tris pH 7.4, 150 mM KCl, 2 mM DTT, 25 mM BSA, 25 mM Salmon Sperm DNA, 5% Glycerol. Enzyme and substrate mixtures were incubated for 30 minutes at 30 °C and separated using 6% Tris-Glycine PAGE gels (Thermo Scientific XP00062BOX). Gels were imaged using a iBright FL1500 gel imager (Thermo Scientific AF44115) and densitometry was performed using iBright Analysis Software (Thermo Scientific).

Activity assays were carried out with 20 nM of each substrate in a buffer consisting of 25 mM Tris pH 8.0, 100 mM KCl, 50 mM NaCl, 1 mM Spermidine, 5 mM MgCl_2_, 2.5 mM DTT, 5% Glycerol. Enzyme and substrate mixtures were incubated for 1 hour at 30 °C and reactions were stopped by addition of Proteinase K and incubation at 37 °C for 30 minutes. Samples were visualized using a 4200 Tapestation (Agilent G2991BA) using D1000 DNA Screen Tapes (Agilent, 5067-5582). Area under the curve measurements for the resulting bands were used to calculate the fraction of each substrate/product. Specific activity calculations were performed using the concentration of enzyme where the reaction had proceeded to 20% completeness.

### *in vitro* transcription of mRNA

All studied genes were cloned into the vector backbone for *in vitro* transcription (IVT) that contains a single copy of the 5’ UTR and the 3’ UTR from the *Xenopus laevis* beta globin gene, in addition to a 110nt polyA tail. Plasmid DNA containing coding sequences were linearized using a BspQI restriction site located immediately downstream of the polyA tail. Linearized plasmids were then purified via AMPure XP magnetic bead (Beckman Coulter A63882).

All mRNAs were generated via IVT reactions using the T7 RNA polymerase. mRNA was capped co-transcriptionally with CleanCap Reagent AG (TriLink BioTech N-7113-100). Each reaction contained final concentration of ∼50 µg/mL for linearized DNA template, 1X reaction buffer (Hongene Biotech ON-062), 15 mM of MgCl_2_, 10 units/µL of T7 RNA polymerase (Hongene Biotech ON-004), 0.002 units/µL of yeast inorganic pyrophosphatase (Hongene Biotech ON-025), 1 unit/µL of murine ribonuclease inhibitor (Hongene Biotech ON-039), 5 mM of CleanCap Reagent AG (TriLink BioTech N-7113-100), and 5 mM of each NTP (Jena Bioscience NU-1010, Hongene Biotech R3-056, Hongene Biotech R2-057). In each IVT reaction, UTP was swapped for N1-methylpseudouridine-5’-triphosphate (BOC Sciences 1428903-59-6). IVT reactions were incubated at 37°C for 2 hours, followed by DNase I digestion of the template DNA (Hongene Biotech ON-109). mRNA products were purified using an RNA Maxi Prep kit (Qiagen), quantified using a NanoDrop (Thermo Fisher Scientific), and checked for integrity using a Fragment Analyzer (Agilent).

### Generation of attB+ HEK293FT cell line

In-house produced lentivirus containing a transfer plasmid with an EF1α-PuroR-WPRE backbone carrying a 46 bp Bxb1 attB insert was transduced into HEK293FT cells (Thermo Fisher Scientific R70007). Cells with Low MOI were plated in sterile 96-well tissue culture plates under puromycin selection via serial dilutions for clone selection. After clone selection, single lentiviral insertion on chromosome 5 was confirmed using ligation mediated PCR with primers targeting the 5’ and 3’ LTRs along with Cergentis TLA mapping (data not shown).

### General cell culture conditions

The attB+ HEK293FT cells were cultured in DMEM (Corning 10-013-CV) with 10% FBS (Gibco A3160501) and 1 ug/mL puromycin under standard tissue culture conditions (at 37°C and 5% CO2).

Cells were detached for splitting and plating using TrypLE Express (Thermo Fisher Scientific 12604039) according to the manufacturer’s instructions.

### Transfection conditions

To transfect cells with plasmids and mRNAs, Lipofectamine MessengerMax (Thermofisher LMRNA001) was used as transfection reagent. 1 d before transfection, cells were seeded onto 96-well tissue culture plates at a density of 21,000 cells per well. For the on-beacon integration assay, 30 ng of Bxb1 variant mRNA and 3.5 ng of cargo plasmid were mixed with 0.3 uL of Lipofectamine MessengerMax and co-transfected into cells plated in a single well. For the off-beacon integration assay, 100 ng of Bxb1 variant mRNA and 60 ng of cargo plasmid were mixed with 0.3 uL of Lipofectamine MessengerMax and co-transfected into cells plated in a single well.

### Genomic DNA extraction

Genomic DNA (gDNA) was extracted 3 d after transfection by removing medium, resuspending cells in 50 uL QuickExtract (LGC Biosearch Technologies QE0905T), and incubating at 75°C for 10 min followed by 95°C for 5 min. Then gDNA was purified from cell lysates by AMPure XP magnetic bead (Beckman Coulter A63882).

### Droplet digital polymerase chain reaction (ddPCR)

Custom primers and probes were designed to measure editing in the studied loci. Probes were dual labelled with 3’-3IABkFQ and either 5’-carboxyflurescein (FAM) for edit targets or 5’-hexachloro-fluorescein phosphoramidite (HEX) for reference genes. From each sample, ddPCR signals from the edit target assay and reference assay were collected together. Percent integration was calculated by dividing the number of FAM-positive droplets (indicating successful cargo integration) by the total number of HEX-positive droplets (reference gene) and then converting the values to a percentage. Assays were validated using gBlocks representing edit outcomes to test for both specificity and linearity. All primers, probes, and gBlocks were synthesized by Integrated DNA Technologies (IDT).

Each reaction contained 11 µL of 2x ddPCR Supermix for probes (No dUTP) (Bio-Rad 1863025), final concentration of 0.5 µM for each primer and 0.25 µM for each probe, 0.11 µL each of HindIII and Eco91I (Thermo Fisher Scientific FD0504 and FD0394, respectively) ∼100 ng of gDNA and water to a final volume of 22 µL. Droplets were generated on the AutoDG Instrument for automated droplet generation (Bio-Rad 1864101). PCR amplification was performed with the following cycling parameters: initial enzyme activation at 94°C for 10 min, followed by 40 cycles of denaturation at 94°C for 30 s and combined annealing/extension step at 58°C for 1 min, and a final step at 98°C for 5 min. Data acquisition and analysis were performed on the QX200 Droplet Reader (Bio-rad 1864003).

### In vivo mouse studies

All animal study procedures were approved by Explora BioLabs under IACUC protocol EB17-004-302. Transgenic C57BL/6J mice with a knock-in of the attB site in intron 1 of F9 or Rosa26 were generated by Biocytogen (Beijing, China) using CRISPR/Cas9. Mice were transferred to Biomere (Worcester, MA, USA) for breeding. AAV8 (AAVG107) cargo was intravenously injected into adult mice (∼20-46 weeks) at a dose of 2E13 or 2.5E13 at day zero. At day seven, LNPs formulated with corresponding Bxb1 variant mRNAs were intravenously injected into the mice at a dose of 3 mg/kg. On day fourteen post LNP injection, animals were euthanized, and liver tissue was collected from the median lobe from each animal and homogenized on Precellys Evolution (cat K002198-PEVO0-A.0 Combo, Bertin technologies, WA, USA).

### Liver gDNA isolation and analysis

Liver gDNA was extracted with quick-DNA/RNA MagBead kit (cat R2131, Zymo research, CA, USA) and analyzed by ddPCR using BioRad Automated Droplet Generator (cat 1864101) and BioRad QX200 Droplet Reader (cat 1864003). Approximately 30 ng of input gDNA were loaded into all ddPCR reactions. Restriction enzymes Eco911 and HindIII were added at [1:200] dilution to ddPCR master mixes. I-PGI attL and attR analyses were performed using the following ddPCR primer-probes: **mF9 attB** (*attL*: Fwd primer (5’-GGAGTCCAACCTTAAGTCTCTTG-3’), Rev primer (5’-GGTTCAGGATCTTGTTGGCG-3’), probe (5’-FAM CTTGTCGACGACGGCGGTCTCAGTG-3’); *attR*: Fwd primer (5’-GCAACTTGGGCTACCAGAT-3’), Rev primer (5’-GCTGTTTTGTTTCCACTGTGTAC-3’), probe (5’-FAM CGGTCTCCGTCGTCAGGATCAT-3’), **Rosa26 attB**: (*attL*: Fwd primer (5’-AGTATACCAGCTAGCAGGCC-3’), Rev primer (5’-GGTTCAGGATCTTGTTGGCG-3’), probe (5’-FAM CTTGTCGACGACGGCGGTCTCAGTG-3’); *attR*: Fwd primer (5’-GCAACTTGGGCTACCAGAT-3’), Rev primer (5’-GCAGAGAACTCCCAGAAAGGT-3’), probe (5’-FAM CGGTCTCCGTCGTCAGGATCAT-3’). mTfrc probe (Applied Biosciences cat 4458370) was used as reference.

